# Evidence-Based Design and Evaluation of a Whole Genome Sequencing Clinical Report for the Reference Microbiology Laboratory

**DOI:** 10.1101/199570

**Authors:** Anamaria Crisan, Geoffrey McKee, Tamara Munzner, Jennifer L. Gardy

**Affiliations:** Dept. of Computer Science, University of British Columbia, Vancouver, BC, CANADA; School of Population and Public Health, University of British Columbia, Vancouver, BC, CANADA; British Columbia Centre for Disease Control, Vancouver, BC, CANADA

## Abstract

**Background:** Microbial genome sequencing is now being routinely used in many clinical and public health laboratories. Understanding how to report complex genomic test results to stakeholders who may have varying familiarity with genomics – including clinicians, laboratorians, epidemiologists, and researchers – is critical to the successful and sustainable implementation of this new technology; however, there are no evidence-based guidelines for designing such a report in the pathogen genomics domain. Here, we describe an iterative, human-centered approach to creating a report template for communicating tuberculosis (TB) genomic test results.

**Methods:** We used Design Study Methodology – a human centered multi-stage approach drawn from the information visualization domain – to redesign an existing clinical report. We used expert consults and an online questionnaire to discover various stakeholders’ needs around the types of data and tasks related to TB that they encounter in their daily workflow. We also evaluated their perceptions of and familiarity with genomic data, as well as its utility at various clinical decision points. These data shaped the design of multiple prototype reports that were compared against the existing report through a second online survey, with the resulting qualitative and quantitative data informing the final, redesigned, report.

**Results:** We recruited 78 participants, 65 of whom were clinicians, nurses, laboratorians, researchers, and epidemiologists involved in TB diagnosis, treatment, and/or surveillance. Our first survey indicated that participants were largely enthusiastic about genomic data, with the majority agreeing on its utility for certain TB diagnosis and treatment tasks and many reporting some confidence in their ability to interpret this type of data (between 58.8% and 94.1%, depending on the specific data type). When we compared our four prototype reports against the existing design, we found that for the majority (86.7%) of design comparisons, participants preferred the alternative prototype designs over the existing version, and that both clinicians and non-clinicians expressed similar design preferences. Participants articulated clearer design preferences when asked to compare individual design elements versus entire reports. Both the quantitative and qualitative data informed the design of a revised report, which is available online as a LaTeX template.

**Conclusions:** We show how a human-centered design approach integrating quantitative and qualitative feedback can be used to design an alternative report for representing complex microbial genomic data. We suggest experimental and design guidelines to inform future design studies in the bioinformatics and microbial genomics domains, and suggest that this type of mixed-methods study is important to facilitate the successful translation of pathogen genomics in the clinic, not only for clinical reports but also more complex bioinformatics data visualization software.

## INTRODUCTION

Whole Genome Sequencing (WGS) is quickly moving from proof-of-concept research into routine clinical and public health use. WGS can diagnose infections at least as accurately as current protocols Fukui et al. (2015); Loman et al. (2013), can predict antimicrobial resistance phenotypes for certain drugs Bradley et al. (2015); Pankhurst et al. (2016); Walker et al. (2015) with high concordance to culture-based testing methods, and can be used in outbreak surveillance to resolve transmission clusters at a resolution not possible with existing genomic or epidemiological methods Nikolayevskyy et al. (2016). Importantly, WGS offers faster turnaround times compared to many culture-based tests, particularly for antimicrobial resistance testing in slow-growing bacteria.

As reference microbiology laboratories move towards accreditation of WGS for routine clinical use, the community is turning its attention toward standardization – developing standard operating procedures for reproducible sample handling, sequencing, and downstream bioinformatics analysis Budowle et al. (2014); Gargis et al. (2016). Reporting genomic microbiology test results in a way that is interpretable by clinicians, nurses, laboratory staff, researchers, and surveillance experts and that meets regulatory requirements is equally important; however, relatively little effort has been directed toward this area. WGS clinical reports are often produced in-house on an *ad hoc*, project-by-project basis, with the resulting product not necessarily meeting the needs of the many stakeholders using the report in their clinical and surveillance workflows.

### Human-Centered Design in the Clinical Laboratory

The information visualization, human-computer interaction, and usability engineering fields offer techniques and design guidelines that have informed bioinformatics tools, including Disease View Driscoll et al. (2011) for exploring host-pathogen interaction data and Microreact Argimón et al. (2016) for visualizing phylogenetic trees in the context of epidemiological or clinical data. Although the public health community is beginning to recognize the potential role of visualization and analytics in daily laboratory workflows Carroll et al. (2014) these techniques have not yet been applied to routine reporting of microbiological test results. However, work from the human health domain – particularly the formatting and display of pathology reports, where standardization is critical Leslie and Rosai (1994) – sheds light on the complex task of clinical report design.

Valenstein reports four principles for organizing an effective pathology report: use headlines to emphasize key points, ensure design continuity over time and relative to other reports, consider information density, and reduce clutter Valenstein (2008), while Renshaw *et al.* note that when pathology report templates were reformatted with numbering and bolding to highlight required information, template completion rates rose from 84 to 98% Renshaw et al. (2014). Fixed, consistent layout of medical record elements, highlighting of data relative to background text, and single-page layout improve clinicians’ ability to locate information Nygren et al. (1998), while information design principles, including visually structuring the document to separate different elements and organizing information to meet the needs of multiple stakeholder types, can reduce the number of errors in data interpretation Wright et al. (1998).

Work in the electronic health record (EHR) and patient risk communication domains has also provided insight into not just the final product but also the process of effective design. Through quantitative and qualitative evaluations, research has shown that some EHRs are difficult to use because they were not designed to support clinical tasks and information retrieval, but rather data entry Wright et al. (1998). Reviews of the risk communication literature note that, while many visual aids improve patients’ understanding of risk Zipkin et al. (2014), the design features that viewers preferred – namely simplistic, minimalist designs – were not necessarily those that led to an accurate interpretation of the underlying data Ancker et al. (2006). Together, these gaps indicate a need for a human-centered, participatory approach iteratively incorporating both design and evaluation Hettinger et al. (2017); Horsky et al. (2012).

### Collaboration Context – COMPASS-TB

The COMPASS-TB project was a proof-of-concept study demonstrating the feasibility and utility of WGS for diagnosing tuberculosis (TB) infection, evaluating an isolate’s antimicrobial sensitivity/resistance, and genotyping the isolate to identify epidemiologically related cases Pankhurst et al. (2016). On the basis of COMPASS-TB’s results, Public Health England (PHE) has implemented routine WGS in the TB reference laboratory PHE (2016); however, this requires changing how mycobacteriology results are reported to clinical and public health stakeholders. The COMPASS-TB pilot used reports designed by the project team, but as clinical implementation within PHE progressed, team members expressed an interest in redesigning the report (Figure 1) to facilitate interpretation of this new data type and align laboratory reporting practices with the needs of multiple TB stakeholders.

**Figure 1.**
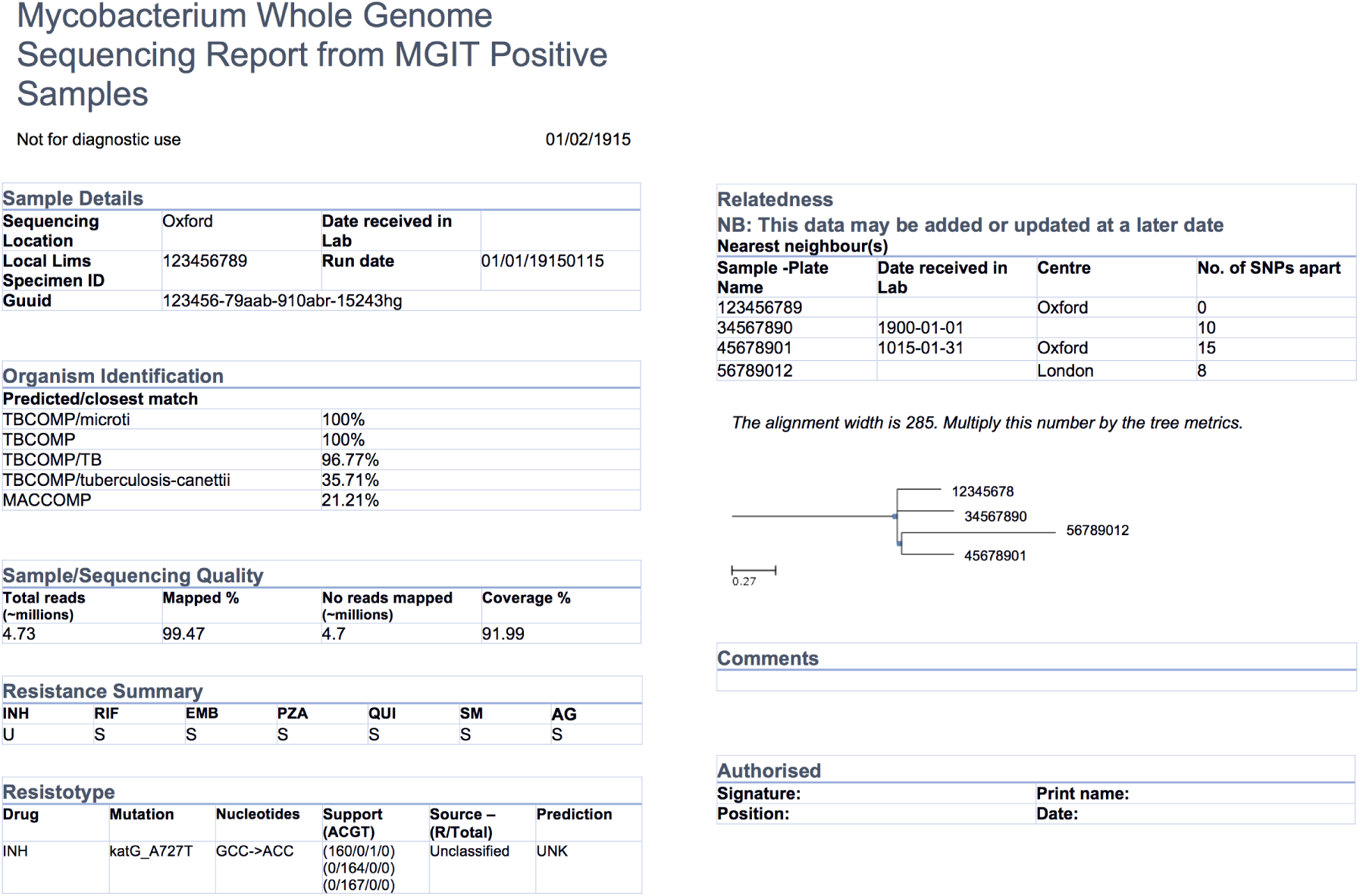
An earlier COMPASS-TB report design.

We undertook a mixed-methods and iterative human-centered approach to inform the design and evaluation of a clinical TB WGS report. Specifically, we chose to use Design Study Methodology Sedlmair et al. (2012) – an approach adopted from the information visualization discipline. When using a Design Study Methodology approach, researchers examine a problem faced by a group of domain specialists, explore their available data and tasks they perform in reference to that problem, create a product (in our case a report, but in the more general case a visualization system) to help solve the problem, assess the product with domain specialists, and reflect on the process to improve future design activities. Compared to an *ad hoc* approach to design, Design Study Methodology engages domain specialists and grounds the design and evaluation of the visualization system in tasks – in this case TB diagnosis, treatment, and surveillance – as well as data. It is this marriage of data and tasks to design choices informed by real needs and supported by empirical evidence that results in a final product that is relevant, usable, and interpretable.

Here we describe our application of design study methodology to the COMPASS-TB report redesign. We show how evidence-based design can be incorporated into the emerging field of clinical microbial genomics, and present a final report template, which may be ported to other organisms. We also recommend a set of guidelines to support future applications of human-centered design in microbial genomics, whether for report designs or for more complex bioinformatics visualization software.

## MATERIALS AND METHODS

### Overview of Design Study Methodology

The Design Study Methodology Sedlmair et al. (2012) is an iterative framework outlining an approach to human-centered visualization design and evaluation. It consists of three phases – Precondition, Core Analysis, and Reflection – that together comprise nine stages. The Precondition and Reflection phases focus on establishing collaborations and writing up research findings, respectively, and are not elaborated upon further here. We describe our work within each of the three stages of the Core Analysis phase: Discovery, Design, and Implementation (Figure 2). We define *domain specialists* in this case as the TB stakeholders — clinicians, laboratorians, and epidemiologists -– who regularly use reports from the reference mycobacteriology laboratory in their work.

**Figure 2.**
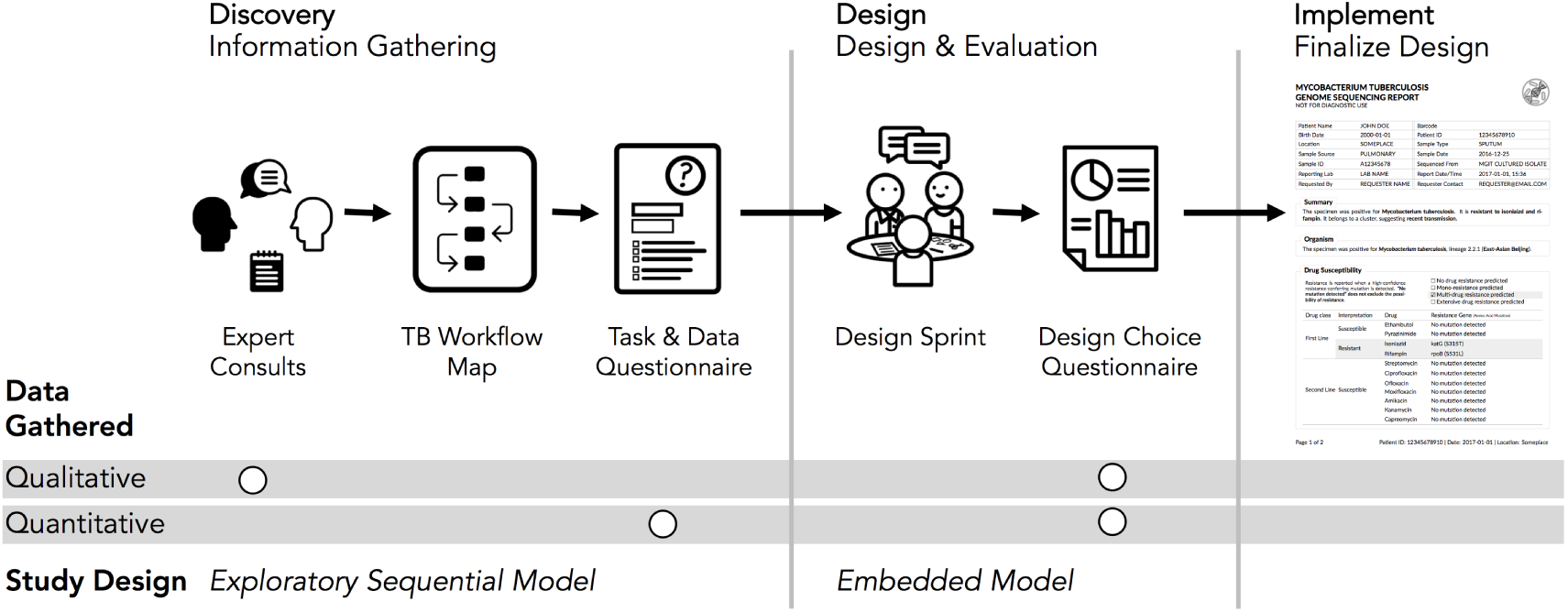
Our human-centered design approach. The Core Analysis phase of the Design Study Methodology consists of Discovery, Design, and Implementation stages. Using this methodological backbone, we collected and analyzed data using mixed-methods study designs in the Discovery and Design stages, which informed the final TB WGS clinical report design.

Our research was reviewed and approved by the University of British Columbia’s Behavioural Research Ethics Board (H10-03336). All data were collected through secure means approved by the university and were de-identified for analysis and sharing. Anonymized *quantitative* results from each of the surveys and the analysis code are available at https://github.com/amcrisan/TBReportRedesign and in the supplemental materials. We also provide the full text of our survey instruments in the Supplemental Materials.

### Discovery Stage

In the Discovery stage, we used an exploratory sequential model Creswell (2014), first gathering qualitative data through expert consults to identify the data types used in TB diagnosis, treatment, and surveillance tasks and then gathering quantitative data through an online survey to more robustly link particular data types to specific tasks.

Our expert consults took the form of semi-structured interviews with seven individuals recruited from the COMPASS-TB project team, the British Columbia Centre for Disease Control (BCCDC), and the British Columbia Public Health Laboratory (BCPHL). The interview questions served as prompts to structure the conversation, but experts were free to comment, at any depth, on the different aspects of TB diagnosis, treatment, and surveillance. We took notes during the consults in order to identify the tasks and data types common to TB workflows in the UK and Canada, as well as to determine which tasks could be supported by WGS data.

Informed by the expert consults, we drafted a Task and Data Questionnaire (text in Supplemental Materials) to survey data types used across the TB workflow (see Figure 3 for a list of data types), the role for WGS data in diagnosis, treatment, and surveillance tasks, and participants’ confidence in interpreting different data types. The questionnaire primarily used multiple choice and true/false type questions, bu also included the optional entry of freeform text. The questionnaire was deployed online using the FluidSurveys platform and participants were recruited using snowball and convenience sampling for a one-week period in July, 2016. For questions pertaining to diagnostic and treatment tasks, we gathered information only from participants self-identifying as clinicians; for the remaining sections of the survey, all participants were prompted to answer each question.

**Figure 3.**
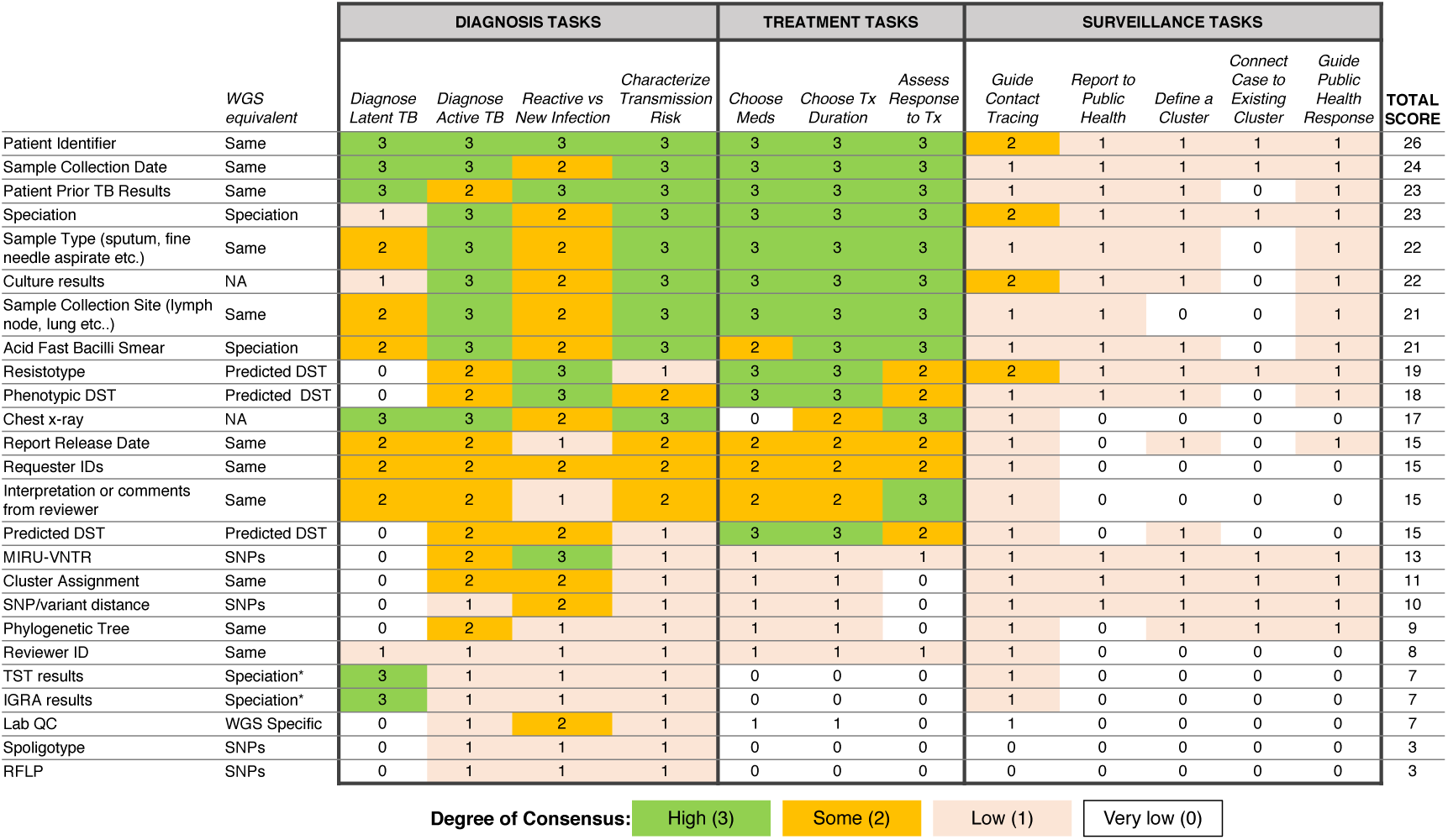
Extent of consensus between TB workflow tasks and available TB data.

Only completed questionnaires were used for analysis. For questions pertaining to participants’ background, their perception of WGS utility, and their confidence interpreting WGS data, we report primarily descriptive statistics. To link TB workflow tasks to specific data types, we presented participants with different task-based scenarios related to diagnosis, treatment, and surveillance and asked which data types they would use to complete the task. For each pair of data and task we assigned a consensus score depending on the proportion of participants who reported using a data type for a specific task: 0 for fewer than 25% of participants, 1 for 25-50%, 2 for 50-75%, and 3 if more than 75% of participants reported using a specific data type for the task at hand. Consensus scores for a data type were also summed across the different tasks. Freeform text, when it was provided, was considered only to add context to participant responses.

### Design Stage

The Discovery stage revealed which data types to include in the redesigned report, while the goal of the Design stage was to identify how it should be presented. We used a Design Sprint event to produce a series of prototype reports, which were then assessed through a second online questionnaire. Using an embedded mixed methods design Creswell (2014), this survey collected quantitative and optional qualitative data on participants’ preference for specific design elements.

The Design Sprint was an interactive design session involving members of the University of British Columbia’s Information Visualization research group, in which teams created alternative designs to report WGS data for the diagnosis, treatment, and surveillance tasks. Teams developed paper prototypes Lloyd and Dykes (2011); Vredenburg et al. (2002) of a complete WGS TB report and, at the completion of the event, presented their prototypes and the rationale for each design choice. The paper prototypes were then digitally mocked up, both as complete reports and as individual elements (see the results in Figure 4 and Figure 5); these digital prototypes were standardized with respect to text, fonts, and sample data where appropriate and used as the basis of the second online survey.

**Figure 4.**
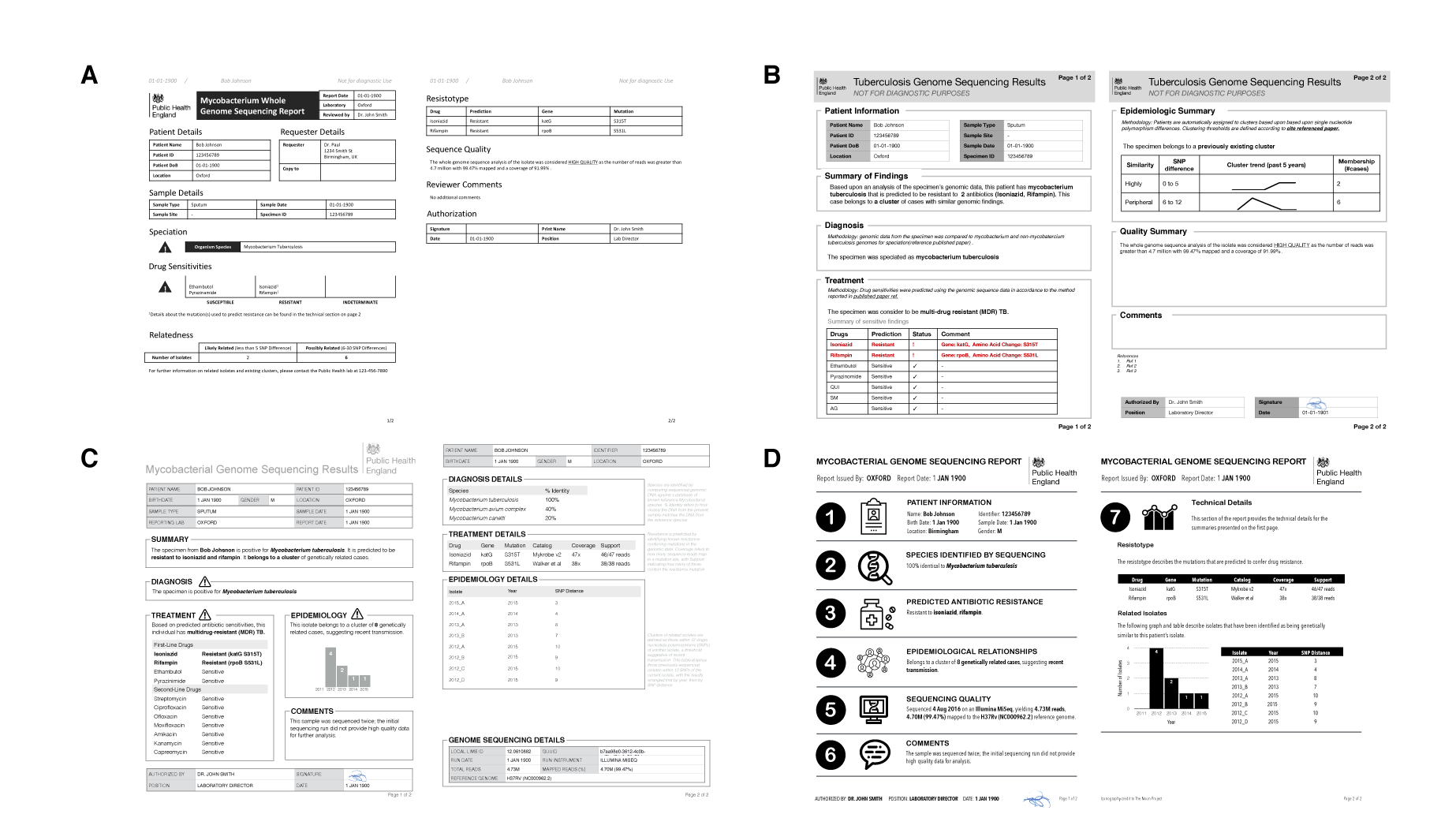
Digital mockups of complete report prototypes generated during the design sprint.

**Figure 5.**
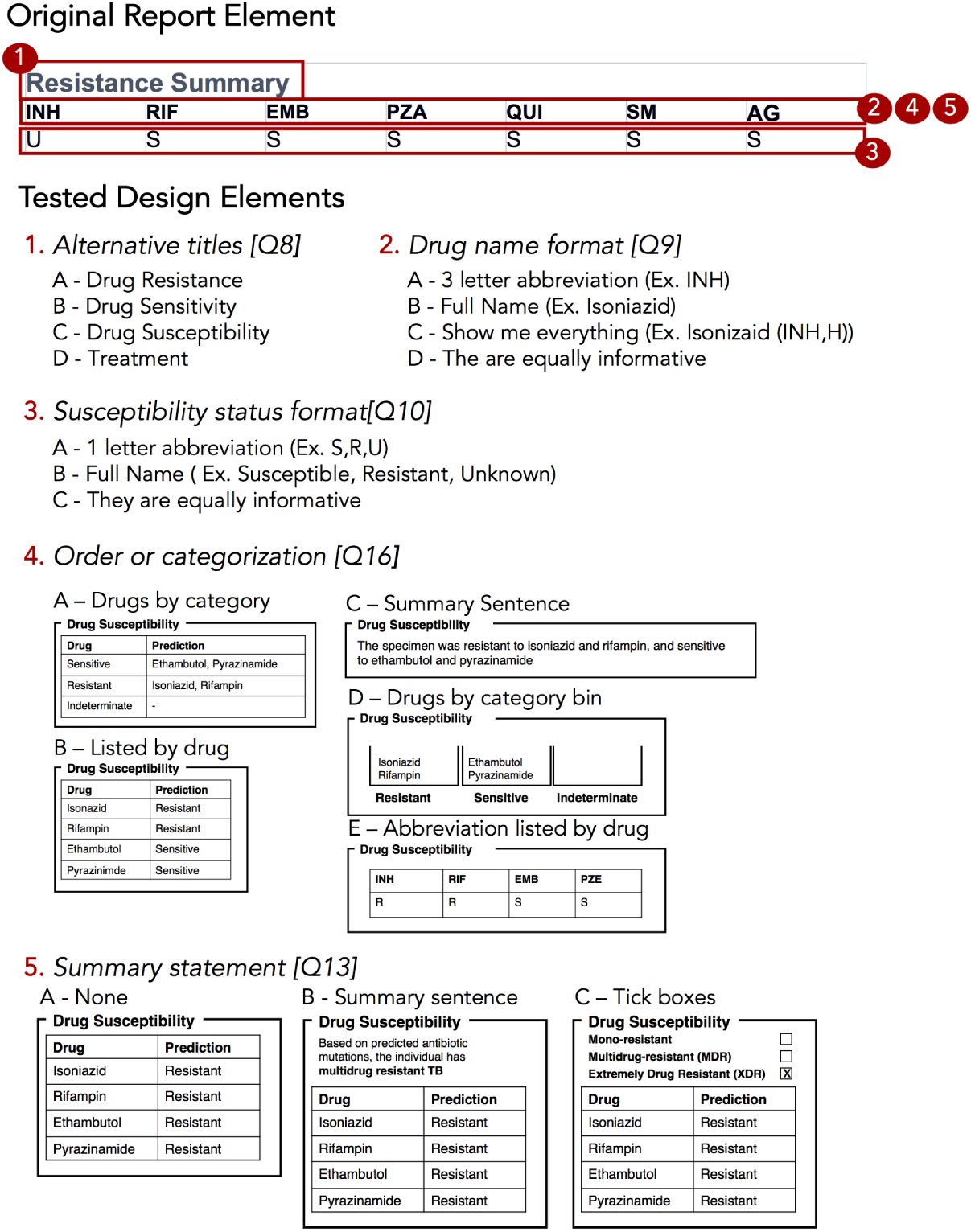
Isolated design elements. The original report element, highlighted in red, is broken down into isolated design elements, each of which was tested independently in the report design survey. In this example, the original resistance summary yields five different alternative wordings and design elements.

In the Design Choice Questionnaire (text in Supplemental Materials), we evaluated participants’ preferences for individual design elements, comparing the options generated during the Design Sprint as well as the initial COMPASS-TB report design, which we hereafter refer to as the control design. As with the first survey, the questionnaire used FluidSurveys, with participants recruited using snowball and convenience sampling. Individuals who had previously participated in the Data and Task Questionnaire were also invited to participate. The survey was open for one month beginning September 10, 2016 and was reopened to recruit additional participants for one month beginning January 5, 2017, as part of the registration for a TB WGS conference hosted by PHE. Only completed surveys were analyzed.

We used single-selection multiple-choice, Likert scale, and ranking questions to assess participant preferences. For multiple-choice and Likert scale questions, we calculated the number of participants that selected each option and report the sum. For questions that required participants to rank options we calculated a rescaled rank score as follows:

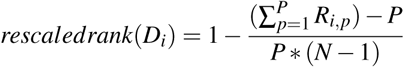

where for each design choice (*D*_*i*_), *i* = {1…*N*} and N is the total number of design choices, *R* = {1…*N*} is a raw rank (rank selected by a participant in the study), and *P* = {1…*P*} is the total number of participants. In our study, 1 was the highest rank (most preferred) and N was the lowest rank (least preferred) option. As an example, if a some design, *D*_1_, is always ranked 1 (greatest preference by everyone), the sum of those ranks is P, resulting in a numerator of 0, and a rescaled rank score of 1; alternatively, if a design, *D*_2_, is always ranked last (N), the sum of those ranks will be P*N, which results in a numerator of P * (N-1), and a rescaled rank score of 0. Thus, the rescaled rank score ranges from 1 (consistently ranked as first) to 0 (consistently ranked last). This transformation from raw to rescaled ranks allows us to compare across questions with different numbers of options, but is predicated on each design alternative having a rank, which is why this approach was not extended to multiple choice questions.

To contextualize rescaled rank scores, we randomly permuted participants’ scores 1000 times and pooled the rescaled rank scores across these iterations to obtain an average score (intuitively and empirically this is 0.5 for the rank questions and 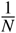 for multiple choice questions) and standard deviation. For each design choice, we plotted its actual rescaled rank score against the distribution of random permutations, highlighting whether the score was within ±1, 2, or 3 standard deviations from the random permutation mean score. The closer a score was to the mean, the more probable that the participants’ preferences were no better than random.

### Implementation Stage

By combining the results of the Design Choice Questionnaire with medical test reporting requirements from the ISO15189:2012 standards, we developed a final template for reporting TB WGS data in the clinical laboratory. The final prototype is implemented in Latex and is available online as a template accessible at: http://www.cs.ubc.ca/labs/imager/tr/2017/MicroReportDesign/.

## RESULTS

Expert consults, the Task and Data Questionnaire, and the Design Choice Questionnaires recruited a total of 78 participants across different roles in TB management and control (Table 1).

**Table 1.**
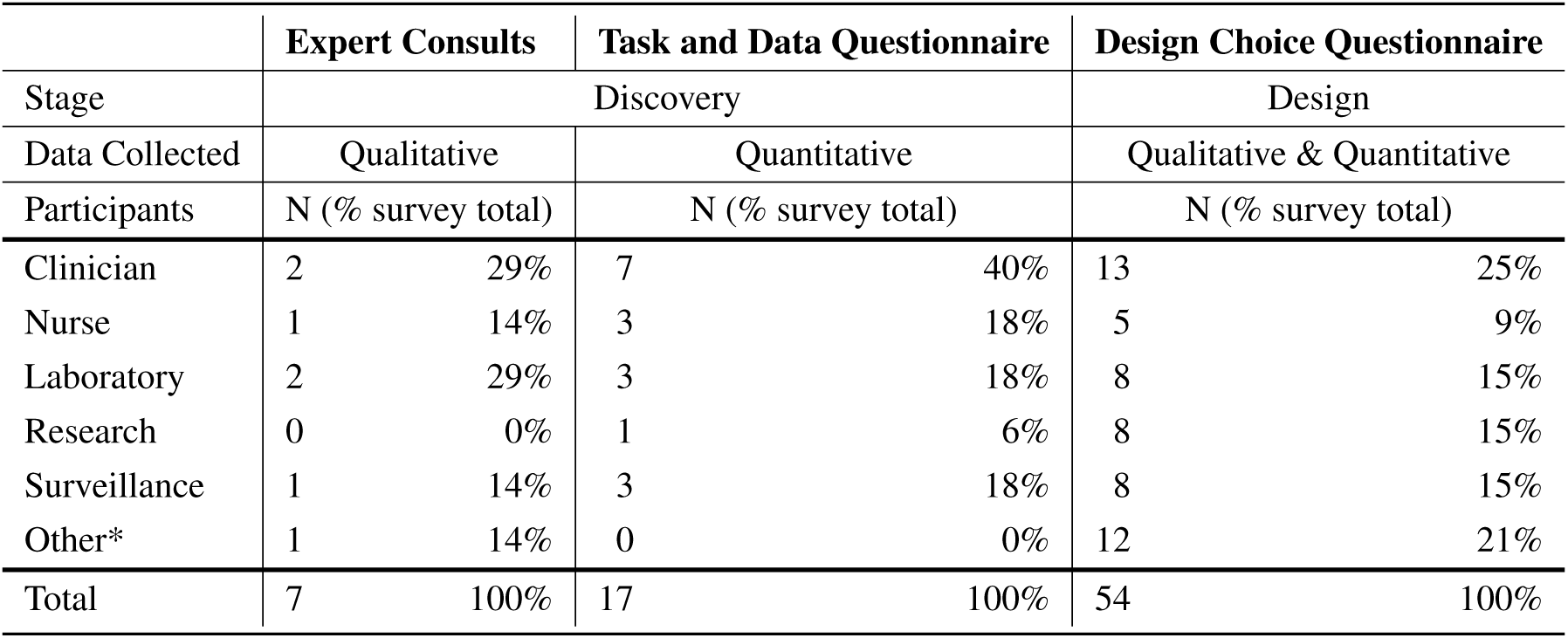
Total study participants across different stages of the Design Study Methodology.

### Experts Emphasized Prioritizing Information and Revealed Constraints

The objective of our expert consults was to understand how reports from the reference mycobacteriology laboratory are currently used in the day-to-day workflows of various TB stakeholders, including clinicians, laboratorians, epidemiologists, and researchers, and what data types are currently used to inform those tasks. Tasks and data types enumerated in the interviews were used to populate downstream quantitative questionnaires; however, the interviews also provided insights into how stakeholders viewed the role of genomics in a clinical laboratory.

Amongst the procedural insights, stakeholders frequently reported that the biggest benefit of WGS over standard mycobacteriology laboratory protocols was to improve testing turnaround times and gather all test results into a single document, rather than having multiple lab reports arriving over weeks to months. Several experts emphasized that these benefits can only be realized if the WGS analytical pipeline has been clinically validated. Although our study team included a clinician and a TB researcher, two surprising procedural insights emerged from the consultations. First, multiple experts from a clinical background emphasized that this audience has extremely limited time to digest the information found on a clinical report. In describing their interaction with a laboratory report, one participant noted that *“10 seconds [to review content] is likely, one minute is luxurious”* while others described variations on the theme of wanting bottom-line, actionable information as quickly as possible. This insight profoundly shaped downstream decisions around how much data to include on a redesigned report and how to arrange it over the report to permit both a quick glance and a deeper dive. Second, experts indicated that laboratory reports were delivered using a variety of formats, including PDFs appended to electronic health records, faxes, or physical mail. This created design constraints at the outset of the project – our redesigned report needed to be legible no matter the medium, ruling out online interactivity, and needed to be black and white.

### Experts Vary in Their Perception of Different Data Types

At the data level, we observed that the experts had differing perceptions of data types and desired level of detail between clinicians and non-clinicians, perhaps reflecting the clinicians’ procedural need for rapid interpretation. Clinicians emphasized the importance of presenting actionable results clearly and omitting those that were not clinically relevant for them. For example, when presented with the sequence quality data on the current COMPASS-TB report (Figure 1) – metrics reflecting the quality of the sequencing run and downstream bioinformatics analysis – interviewees did not expect the lab to release poor quality data, given the presence of strict quality control mechanisms. ISO15189:2012 standards require some degree of reporting around the measurement procedure and results, but this insight suggested such data might best be placed later in the report, in a very simplified format, after the actionable data, or described in the report comments. Similarly, experts were also divided on the interpretability and utility of the phylogenetic tree in the epidemiological relatedness section of the current COMPASS-TB report, with clinicians noting that the case belonging to an epidemiological cluster would not impact their use of the genomic test results.

Experts also disagreed about the level of detail needed for WGS data, and this appeared to depend upon on whether the expert was a clinician as well as their prior experience with WGS through the COMPASS-TB project. For example, one expert indicated that *“clinicians are wanting to know which mutations conferred resistance”*, while another noted that they *“don’t use these [mutations] right now routinely, so it’s not that relevant”*. When asked to comment on the resistance summary table in the current COMPASS-TB report (Figure 1), clinicians were concerned about the use of abbreviations for both drug names and susceptibility status leading to misinterpretation, and many were uncertain how to use the detailed mutation information in the resistotype table.

### WGS Data is Vital, But Some Lack Confidence in its Interpretation

The expert consults provided a detailed overview of the tasks and data associated with TB care, allowing us to create a draft workflow outlining the TB diagnosis, treatment, and surveillance tasks coupled to the supporting data sources and data types (Fig. S1). This workflow was used to design the Task and Data Questionnaire.

Of the 17 participants responding in full to the Task and Data Questionnaire (Table 1), most were from the United Kingdom (88%) and most reported professional experience and formal education in infectious diseases and epidemiology (Table S1). Participants were less likely to report education at the masters or doctoral level in microbial genomics, biochemistry, or bioinformatics (Table S1). Fewer than half (47.1%) of participants had participated in TB WGS projects, but all (100%) participants were enthusiastic about the role of microbial genomics in infectious disease diagnosis, both today (47.1%) and in the near future, pending clinical validation (52.9%).

When queried about their potential future use of molecular data, whether WGS, genotyping, or other, participants indicated they foresaw themselves consulting, often or all the time, data on resistance-conferring mutations (82.3% of participants), MIRU-VNTR patterns (88.2%), epidemiological cluster membership (76.5%), single nucleotide polymorphism/variant distances from other isolates (64.7%), and WGS quality metrics (58.8%) (Table S2). However, of the 14 different data types queried, the majority of participants only felt confident in interpreting four (MIRU-VNTR, drug susceptibility from culture, drug susceptibility from PCR or LPA, genomic clusters) - most participants only felt somewhat confident, or not confident at all, interpreting the other data types (Table S3).

Moving from confidence in their own interpretation of laboratory data types to confidence in the utility of WGS data in general, the majority of participants were confident that information contained within the TB genome *can* be used to correctly perform organism speciation (76.5%), assign a patient to existing clusters (70.0%), rule out transmission events (64.7%), and to a lesser extent were confident TB WGS could be used to identify epidemiologically related patients (58.8%) and predict drug susceptibility (52.9%) (Table S4). The majority of participants thought genomic data *may* be able to inform clinicians of appropriate treatment regimens (100%) and identify transmission events (94.1%); however, participants showed mixed consensus toward whether genomic data could be used to monitor treatment progress for TB (47.2%) or diagnose active TB (52.9%).

### Respondent Consensus Suggests a Role for WGS in Diagnosis and Treatment Tasks

To examine which data types were being used to support diagnosis, treatment, and surveillance tasks in the workflow, we assigned a numerical score reflecting respondent consensus around each data type-task pair (Figure 3). We found greater consensus around the data types that participants would use in diagnosis and treatment tasks, but little consensus around the data they would use for surveillance tasks, contrasting with participants’ previously stated support for using WGS or other genotyping data for understanding TB epidemiology. Overall, the most frequently used data types included administrative data (patient ID, sample type, collection site, collection date) and results from current laboratory tests (solid or liquid culture, smear status, and speciation), which together were used primarily for diagnosis and treatment. Prior test results from a patient were deemed important; however, the earlier expert consults indicated that such data was difficult to obtain and unlikely to be included in future reports.

We also queried participants’ perceptions of barriers impacting their workflow, with the majority of participants (83.3%) reporting issues with both the timeliness of receiving TB data from the reference laboratory and the distribution of test results across multiple documents (Table S5) – a finding that corroborated the procedural insights from the expert consults.

### Prototyping Via a Design Sprint Produces a Range of Design Alternatives

Equipped with an understanding of how WGS data might be used in the various TB workflow tasks, we embarked on the Design stage of the design study methodology. A Design Sprint event involving study team members and information visualization experts resulted in four prototype report designs (Figure 4) and various isolated design elements (Figure 5). Although each prototype used different design elements for the required data types, when the prototypes were compared at the end of the event, common themes emerged. These included: presenting data in an order informed by the workflow – data related to diagnosis, treatment, then surveillance; placing actionable, high-level on the front page, with additional details on the over page; and using both an overall summary statement at the beginning of the report as well as brief summary statements at the beginning of each section.

To drill down and determine which design elements best communicate the underlying data, we isolated individual design elements (Figure 5) and classified them as wording choices – for example, which heading to use for a given section of the report – or design choices, such as layout, the use of emphasis, and the use of graphics (Table S6).

### The Design Choice Questionnaire Quantifies Participant Preferences for Specific Design Elements

We next developed an online survey, the Design Choice Questionnaire, to assess stakeholders’ preferences for both specific design elements and overall report prototypes. The distribution of public health roles amongst survey participants is presented in Table 1; all but 11 participants (20%) actively worked with TB data. Participants were employed by Academic Institutions (35.2%), Hospitals (24.1%), and Public Health Organizations (33.3%), with only 7.4% of participants being employed in some other sector. The majority of participants were from the UK (59.2%), while 11.1% were from Canada; the remaining 29.7% were drawn from the United States (6.5%), Europe (14.8%), Brazil (2.8%), India (2.8%), and Gambia (2.8%)

We first examined participants’ preference for specific wording and design elements (Figure 6A,B), comparing elements arising from the prototypes to those used in the existing COMPASS-TB report, which acted as a control. Notably, of the 15 wording and design elements queried, in only two cases was the control design preferred over a design arising from one of the prototypes (note that one query did not compare to a control). Furthermore, in 8 out of 15 queries (Q6, Q8, Q9, Q10, Q12, Q17, Q5, Q18) participants showed quite strong preferences, assessed by the top preference being +3 or greater standard deviations from the mean for *both* clinicians and non-clinicians, while 7 out of 15 queries (Q14, Q7, Q13, Q16, Q11, Q15, Q19) showed less strongly defined preferences or some discordance of the top choice between clinicians and non-clinicians.

**Figure 6.**
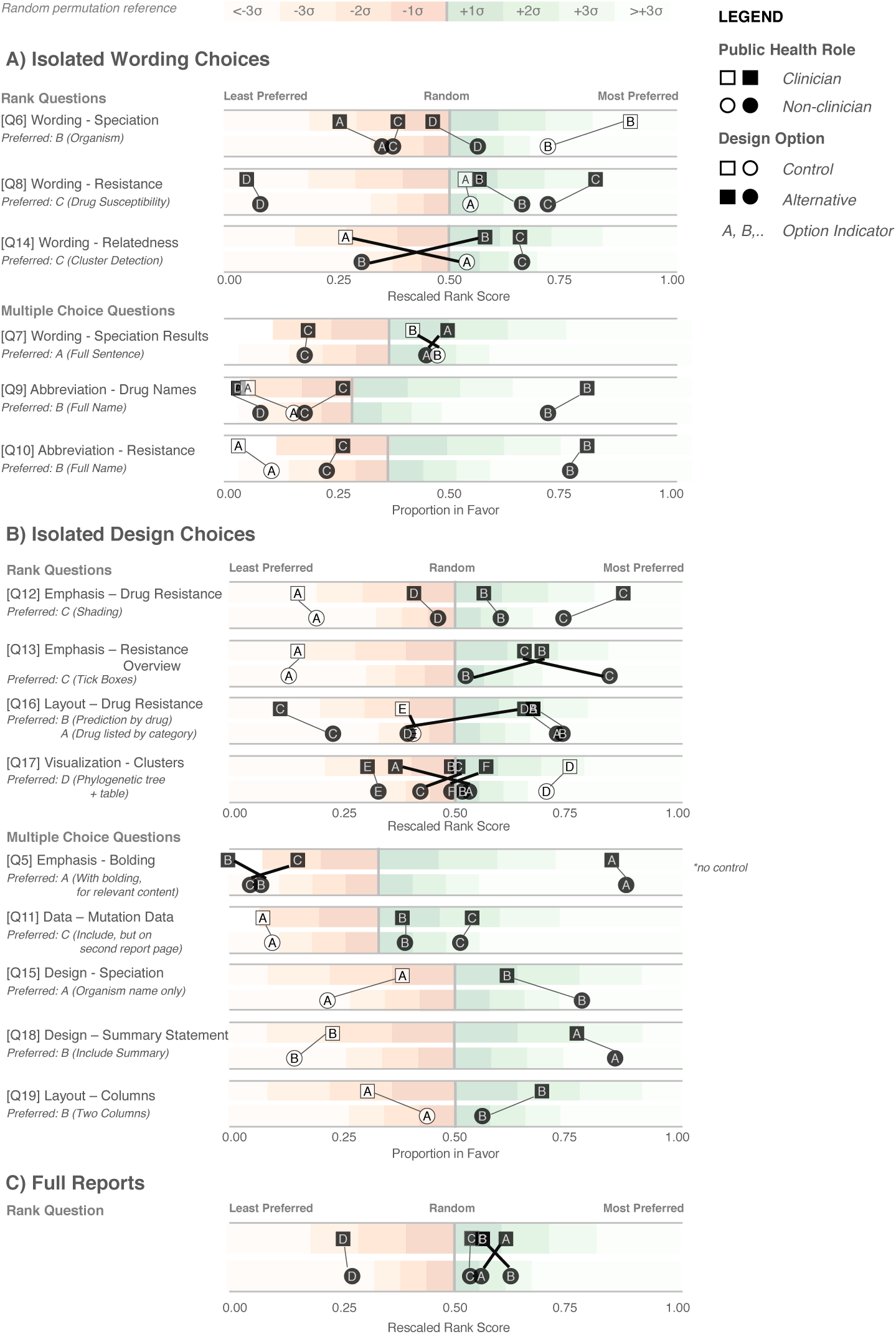
Design Choice Questionnaire results. Responses are grouped according to question type: wording (A), design choices (B), and full reports (C), and partitioned into clinician participants (squares) and non-clinician participants (circles). Responses are colored according to whether they are the control design from the original report (white) or an alternative design devised in the design sprint (black). Lines connect options between clinician and non-clinicians preferences, with thicker crossing lines showing discordance between the two groups and vertical lines showing concordance in preferences. Rescaled rank scores are shown against a reference of random permutations (see Methods), with scores closer to 1 indicating the most preferred response. Specific questions are indicated with Q; the questions as presented to the participants are shown in Table S6

The findings from the analysis of wording elements (Figure 6A) showed that participants preferred complete terms to abbreviations, such as writing out “isoniazid” as opposed to “INH”, or “H” or “resistant” as opposed to “R” and both clinicians and non-clinicians were in agreement over the preferred vocabulary for section headings. Interestingly, wording questions related to the treatment task yielded the widest range of rankings.

Clear preferences were also observed for information design elements, again largely concordant between clinicians and non-clinicians (Figure 6B). Participants preferred elements that drew attention to specific data, such summary statements, shading, and tick boxes, but there was less consensus around how much detail to include and where. The majority of participants indicated that genomic data pertaining to resistance-conferring mutations should be included (Figure 6B; Q11), but were divided as which data should be included and where. Most (85%) wanted to know the gene harboring the resistance mutation (i.e. katG; inhA), but only half wanted details of the specific mutation (50% wanted the amino acid substitution, 46% wanted to know the nucleotide-level change). Many participants preferred that sections be prioritized, with less important details relegated to the second page of the report.

Interestingly, while both clinicians and non-clinicians reported similar rankings for most design elements, one element showed an unusual distribution of scores – the visualization for showing genomic relatedness and membership in a cluster. While both groups of participants preferred a phylogenetic tree accompanied by a summary table, which is the current COMPASS-TB control design, the other four options appeared to be ranked randomly, with rescaled rank score close to 0.5, suggesting that none of the alternative options were particularly good.

We also had participants rank their preferences for the four prototype designs (Figure 6C). While all participants ranked Prototype D as their least preferred choice, many citing that the images used were too distracting, clinicians and non-clinicians varied in their ranking of the other three options, with clinicians preferring option A and non-clinicians preferring B. However, qualitative feedback collected for this question revealed that participants found comparing individual elements easier than comparing full reports.

### Qualitative Data Affords Additional Insights into Report Design

The qualitative responses in the Design Choice Questionnaire raised important points that would otherwise not have been captured by quantitative data alone. For example, the importance of presenting drug susceptibility data clearly emerged from the qualitative responses. Participants indicated that *“the report must call attention [to] drug resistance”* and expressed concern that the abbreviation of drug names and/or predicted resistance phenotype could lead to misinterpretation and pose risks to patient safety, stating that *“not all clinicians [are] likely to recognize the abbreviations”* and *“[using the full name] reduces the risk of errors, especially if new to TB”*. When choosing how to emphasize predicted drug susceptibility information (shading, bolding, alert glyphs, or no emphasis), some participants suggested *“shading draws the quickest attention to [resistance]”* and that *“with presbyopia, resistance can be easily missed and therefore shading affords greater patient safety”*, but other participants indicated drug susceptibility, rather than resistance, should be emphasized: *“not sure that resistant should be shaded – better to shade sensitive drugs in my view”* and *“it would be better to highlight what is working instead of highlight what is not working.”* We opted to highlight resistance given the low incidence of drug-resistant TB in the UK and Canada, which were the primary application contexts. Some reported concerns as to whether such emphasis was possible with current electronic health records, including *“[bolding or shading] may not transfer correctly”* and *“shaded [text] won’t photocopy well”*, which prompted us to test both printing and photocopying of the resulting report.

The issue of clinicians having little time to interact with the report, raised in both the expert consults and the Task and Data Questionnaire, also became apparent in the qualitative responses to the Design Choice Questionnaire, such as “*the best likelihood of success will [come] from the ability to draw attention to someone scanning the document quickly”*. However, participants’ perceptions of which design choices best promoted rapid synthesis varied. Some preferred summaries in the form of check boxes – “*[a] tick box is the most straightforward way to summarize it. Reading a summary sentence will probably take longer”* and *“the check boxes provide an at-a-glance result”* – while others preferred additional commentary – *“interpretation is important; but tick boxes alone lack the necessary nuance required for interpretation”* and that *“tick boxes may cause confusion when clinicians read XDR without realizing that option is not selected. Ideal to add a comment about resistance”.* To address this concern we added a “No drug resistance predicted” option to the check-boxes (absent from the survey design options), and included shading elements to emphasize the drug susceptibility result.

The qualitative responses to Q17 (Figure 6B) provided further insight into the uncertainty around how best to represent genomic relatedness suggestive of an epidemiological relatedness. Some participants felt that data related to surveillance tasks should not appear in a report that is also meant for clinicians, either because it wasn’t relevant to this audience –“*[this data] should not appear in the report. It should only be given to field epi and researchers. Overloading the clinical report would be deteriorating”* and *“not useful for a clinician”* – or because they were uncertain about its interpretation – “*cluster detection would be fine for those who already know what a cluster is”* and “*my patient’s isolate is 6 SNPs from someone diagnosed 3 years ago. What is the clinical action?”.*

Of the design choices for cluster detection, several participants articulated that many of the options, including the control, *“[included] too much information and [were] unnecessary for routine diagnosis/treatment”*. However, others felt that the options did not provide sufficient detail and offered alternatives, such as *“if you can combine the phylogenetic tree with some kind of graph showing temporal spread that would be perfect. Adding geographical data would be a really helpful bonus too.”*. This is an area of reporting that requires further investigation and was not fully resolved in our study.

Finally, participants were candid about those design options that did not work well – for example, of the report design with many graphics (Figure 6A, option D), participants indicated it was *“distracting; looks like a set of roadworks rather than a microbiology report”* and that it was important to *“keep it simple”*. Their feedback also revealed when our phrasing on the survey instruments was unclear.

### Developing a Final Report Template

There are no prescriptive guidelines around integrating our quantitative data, qualitative data, and ISO15189:2012 reporting requirements; thus, we have attempted to be as transparent and empiric as possible in justifying our final design (Figure 7). A more thorough walkthrough is presented in the Supplemental Materials, and here we highlight selected choices. The final prototype is implemented in Latex and is available online as a template accessible at: http://www.cs.ubc.ca/labs/imager/tr/2017/MicroReportDesign/.

**Figure 7.**
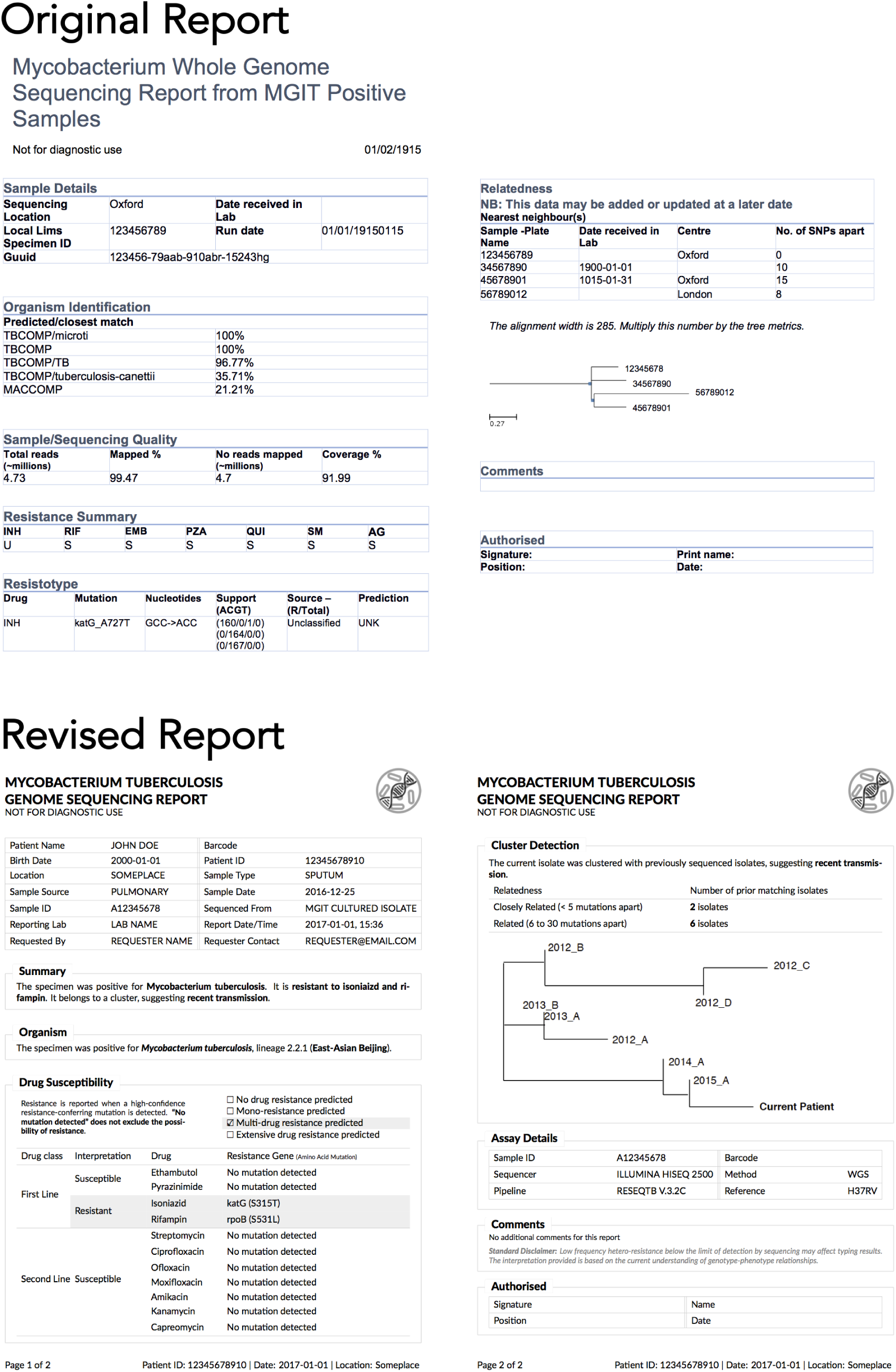
Original and revised reports. The revised report uses empirical evidence gathered through multiple stages of a human centered design process. Note that the image in the upper corner of the revised report is a placeholder for an organizational logo.

We first incorporated ISO15189:2012 requirements (see Supplemental Materials) into the final report template and then turned to the preferences expressed in the Design Choice Questionnaire. Overall, information was structured to mirror the TB workflow – diagnosis, treatment, then surveillance. We chose to limit bolding to relevant information, and used shading to highlight important and actionable clinical information, under the rationale that appropriate use of emphasis could facilitate an accurate and quick reading of the report, with detailed information present but de-emphasized.

In two instances, our design decisions deviated from participant preferences: we opted to use one column instead of two, and we presented detailed genomic resistance data on the first page of the report, rather than the second page. A single column was chosen as all of the information ranked as important by participants could be presented on a single page without the need to condense information into two columns. Because many of the resistotype details of the original report, such as mutation source and individual nucleotide changes (Figure 1), were not included in the revised report, it was possible to present all of the participants’ desired data in a single table on one page.

A draft of the final design was presented to a new cohort of TB stakeholders at a September, 2017 expert working group on standardized reporting of TB genomic resistance data. Through a group discussion, subtle changes to the report were made, including updating some of the language used (for example, replacing occurrences of the word “sensitive” with “susceptible”), adding the lineage to the Organism section, and adding additional fields to tables describing the sample, and the assay, such as what type of material was sequenced (pure culture, direct specimen) and what sequencing platform was used.

## DISCUSSION

Microbial genomics is playing an increasingly important role in public health microbiology, and its successful implementation in the clinic will rely not just on validation and accreditation of WGS-based tests, but also in how effective the resulting reports are to stakeholders, including clinicians. Using Design Study Methodology, we developed a two-page report template to communicate WGS-derived test results related to TB diagnosis, drug susceptibility testing, and clustering.

To our knowledge, this project is the first formal inquiry into human-centered design for microbial genomics reporting. We argue that the application of human-centered design methodologies allowed us to improve not only the visual aesthetics of the final report, but also its functionality, by carefully coupling stakeholder tasks, data, and constraints to techniques from information and graphic design. Giving the original report a “graphic design facelift” would not have improved the functionality, as some of the information in the original report was found to be unnecessary, presented in a way that could lead to misinterpretation,or did not take into account stakeholder constraints. For example, interviews and surveys revealed procedural and data constraints our study team had not anticipated, including the limited time available for clinicians to read laboratory reports and the need for simple, black and white formatting amenable to media ranging from electronic delivery to fax – these findings were critical to shaping the downstream design process. Furthermore, in nearly every case, study participants preferred our alternative design elements, informed by empirical findings in the discovery stage, over the control elements derived from the original report.

Although human-centered information visualization design methodologies are commonly used in software development, it could be asked whether they are warranted in a report design project. One advantage of tackling the simpler problem of report design is that it allows us to demonstrate Design Study Methodology in action and link evidence to design decisions more clearly than with complex software. We also collected data with the intention of applying it to the development and evaluation of more complex reporting and data visualization software that we plan to create. Similarly, others can use our approach or our data to inform the design of simple or complex applications elsewhere in pathogen genomics and bioinformatics.

The exploratory nature of this project brings with it certain limitations. First, our participants were identified through convenience and snowball sampling within the authors’ networks, and thus are likely to be more experienced with the clinical application of microbial genomics. While this is appropriate for the context of our collaboration, in which our goal is redesigning a report for use by the COMPASS-TB team and collaborating laboratories, it does limit our ability to generalize the findings to other settings. Second, we did not have *a priori* knowledge of the effect sizes (i.e. extent of preferential difference for each type of question) in the Design Choice Questionnaire, making sample size calculations challenging. Had *a priori* effect sizes been available, the study could be powered, for example, for the smallest or average effect size. To avoid mis-characterizing our results we have relied on primarily descriptive statistics, without tests for statistical significance, and assert that our findings are best interpreted as first steps toward a better understanding how information *and* visualization design can play a role in reporting pathogen WGS data. Finally, we did not undertake a head-to-head experimental comparison between the original report design and the revised design. While this comparison had been planned at the outset of our project, the results of the Design Choice Questionnaire showed such a clear preference for the alternative designs when comparing isolated components that we concluded there was no need for such a final test as it would yield little new evidence.

For researchers wishing to undertake a similar human-centered design approach, we have summarized our primary findings into three experimental guidelines and five design guidelines. These guidelines arose from our experience throughout this report redesign process, but are intended to apply generally to the process of designing visualizations for microbial genomic data or other human health-related information.

The three experimental guidelines reflect the areas of the design methodology that we found to be particularly important in our data collection and analysis as well as the final report design process. First, **design around tasks**. It is tempting to simply ask stakeholders what they want to see in a final design, but many of them will not be able to create an effective end product because design is not their principal area of expertise. However, stakeholders know very well what they do on a daily basis and can indicate data that are relevant to those specific tasks and can indicate in which areas they require more support. The role of the designer is to marry those tasks, clinical workflows, and constraints into design alternatives. Second, **compare isolated components, and not just whole systems**. Here we use system to mean either a simple report or a more complex software system. Comparing whole systems can overload an individual’s working memory, meaning they may rely on heuristics such as preferences around style or distracting elements, when assessing and comparing full systems Shah and Oppenheimer (2008). Presenting isolated design elements and controlling for non-tested factors (i.e. font, text) can reduce the burden on working memory and isolate the effect of design alternatives. Finally, **compare against a control whenever possible.** If a prior report or system exists, or if there are commonly agreed upon conventions in the literature or field, it is useful to compare novel designs against an existing one. More generally, comparison of multiple alternatives is the most critical defense against defaulting to *ad hoc* designs and the most important step of our human-centered design methodology.

Our five design guidelines reflect techniques from information visualization and graphic design that we used in an attempt to improve the readability of the report and balance different stakeholder information needs. First, **structure information such that it mimics a stakeholder’s workflow**. In this case, the report prioritizes a *clinical* workflow, and this workflow is reflected in the report’s design through the use of gestalt principles Moore and Fitz (1993) to group related data and by ordering information hierarchically so that the document is read according to the clinical narrative we established via feedback from experts and study participants. Second, **use emphasis carefully**. Here, bolding, text size, and shading were reserved to highlight important data and were not applied to aesthetic aspects of the report design. Third, **present dense information in a careful and structured manner.** Stakeholders should not have to search for relevant information – a cognitively expensive task Chang et al. (2012) that can result in information loss Shneiderman (1996). Through the combination of gestalt, visual hierarchy, and careful use of emphasis, it is possible to present a lot of information by creating two layers: a higher-level “quick glance” layer and a more detailed lower layer. The quick glance layer should contain the relevant and clinically actionable information and should be visually salient (i.e “pop-out”), while the detailed layer should be less visually salient and contain additional information that some, but not all, stakeholders may wish to have (based on their tasks and data needs). Fourth, **use words precisely**. Specific terminology may not be uniformly understood or consistently interpreted by stakeholders, particularly when the designer and the stakeholders come from different domains, or even when individuals in the same domain have markedly different daily workflows such as bioinformaticians and clinicians. Finally, **if using images, do so judiciously.** Images can be distracting when they do not convey actionable information relevant to the stakeholder.

## CONCLUSIONS

We applied human-centered design methodologies to redesign a clinical report for a reference microbiology laboratory, but the techniques we used – drawn from more complex applications in information visualization and human-computer interaction – can be used in other scenarios, including the development of more complex data dashboards, data visualization or other bioinformatics tools. By introducing these techniques to the microbial genomics, bioinformatics, and genomic epidemiology communities, we hope to inspire their further use of evidence-based, human-centric design.

## ACKNOWLEDGMENTS

We would like to acknowledge and thank all study participants who took the time to respond to the surveys and provided excellent and valuable insights for our work. We would also like to thank Ana Gibertoni-Cruz, Grace Smith,and Tim Walker for their contributions in the early stages of this project and in recruiting participants, and the members of the Information Visualization group at the University of British Columbia who participated in our Design Sprint: Kimberly Dextras-Romagnino, Dylan Dong, Georges Hattab, and Zipeng Liu. AC is supported by the Vanier Scholars Program, JLG is supported by the Canada Research Chairs Program and the Michael Smith Foundation for Health Research Scholar Award Program, TM is supported by the Natural Sciences and Engineering Research Council of Canada Discovery Program RGPIN-2014-06309. The work described here was funded by the British Columbia Centre for Disease Control Foundation for Population and Public Health, as well as Genome BC, through grant G01SMA: Sharing Mycobacterial Analytic Capacity.

## REFERENCES

Ancker, J. S., Senathrajah, Y., Kukafka, R., and Starren, J. B. (2006). Design features of graphs in health risk communication : a systematic review. Journal of the American Medical Informatics Association, 13(6):608–619.

Argimón, S., Abudahab, K., Goater, R. J. E., Fedosejev, A., Bhai, J., Glasner, C., Feil, E. J., Holden, M. T. G., Yeats, C. A., Grundmann, H., Spratt, B. G., and Aanensen, D. M. (2016). Microreact: visualizing and sharing data for genomic epidemiology and phylogeography. Microbial Genomics.

Bradley, P., Gordon, N. C., Walker, T. M., Dunn, L., Heys, S., Huang, B., Earle, S., Pankhurst, L. J., Anson, L., de Cesare, M., Piazza, P., Votintseva, A. A., Golubchik, T., Wilson, D. J., Wyllie, D. H., Diel, R., Niemann, S., Feuerriegel, S., Kohl, T. A., Ismail, N., Omar, S. V., Smith, E. G., Buck, D., McVean, G., Walker, A. S., Peto, T. E. A., Crook, D. W., and Iqbal, Z. (2015). Rapid antibiotic-resistance predictions from genome sequence data for Staphylococcus aureus and Mycobacterium tuberculosis. Nature Communications, 6:10063.

Budowle, B., Connell, N. D., Bielecka-Oder, A., Colwell, R. R., Corbett, C. R., Fletcher, J., Forsman, M., Kadavy, D. R., Markotic, A., Morse, S. A., Murch, R. S., Sajantila, A., Schmedes, S. E., Ternus, K. L., Turner, S. D., and Minot, S. (2014). Validation of high throughput sequencing and microbial forensics applications. Investigative genetics, 5:9.

Carroll, L. N., Au, A. P., Detwiler, L. T., Fu, T.-C., Painter, I. S., and Abernethy, N. F. (2014). Visualization and analytics tools for infectious disease epidemiology: A systematic review. Journal of Biomedical Informatics, 51:287–298.

Chang, T. W., Kinshuk, Chen, N. S., and Yu, P. T. (2012). The effects of presentation method and information density on visual search ability and working memory load. Computers and Education, 58(2):721–731.

Creswell, J. W. (2014). Research design: qualitative, quantitative, and mixed methods approaches. Thousand Oaks, CA.

Driscoll, T., Gabbard, J. L., Mao, C., Dalay, O., Shukla, M., Freifeld, C. C., Hoen, A. G., Brownstein, J. S., and Sobral, B. W. (2011). Integration and visualization of host-pathogen data related to infectious diseases. Bioinformatics, 27(16):2279–87.

Fukui, Y., Aoki, K., Okuma, S., Sato, T., Ishii, Y., and Tateda, K. (2015). Metagenomic analysis for detecting pathogens in culture-negative infective endocarditis. Journal of Infection and Chemotherapy, 21(12):882–884.

Gargis, A. S., Kalman, L., and Lubin, I. M. (2016). Assuring the quality of next-generation sequencing in clinical microbiology and public health laboratories. Journal of Clinical Microbiology, 54(12):2857–2865.

Hettinger, A. Z., Roth, E. M., and Bisantz, A. M. (2017). Cognitive engineering and health informatics: applications and intersections. Journal of Biomedical Informatics, 67:21–33.

Horsky, J., Schiff, G. D., Johnston, D., Mercincavage, L., Bell, D., and Middleton, B. (2012). Interface design principles for usable decision support: a targeted review of best practices for clinical prescribing interventions. Journal of Biomedical Informatics, 45(6):1202–16.

Leslie, K. O. and Rosai, J. (1994). Standardization of the surgical pathology report: formats, templates, and synoptic reports. Seminars in Diagnostic Pathology, 11(4):253–7.

Lloyd, D. and Dykes, J. (2011). Human-centered approaches in geovisualization design: Investigating multiple methods through a long-term case study. IEEE Transactions on Visualization and Computer Graphics, 17(12):2498–2507.

Loman, N. J., Constantinidou, C., Christner, M., Rohde, H., Chan, J. Z.-M., Quick, J., Weir, J. C., Quince, C., Smith, G. P., Betley, J. R., Aepfelbacher, M., and Pallen, M. J. (2013). A culture-independent sequence-Based metagenomics approach to the investigation of an outbreak of shiga-toxigenic Escherichia coli O104:H4. JAMA, 309(14):1502.

Moore, P. and Fitz, C. (1993). Using Gestalt theory to teach document design and graphics. Technical Communication Quarterly, 2(4):389–410.

Nikolayevskyy, V., Kranzer, K., Niemann, S., and Drobniewski, F. (2016). Whole genome sequencing of Mycobacterium tuberculosis for detection of recent transmission and tracing outbreaks: A systematic review. Tuberculosis, 98:77–85.

Nygren, E., Wyatt, J. C., and Wright, P. (1998). Helping clinicians to find data and avoid delays. Lancet, 352(9138):1462–1466.

Pankhurst, L. J., del Ojo Elias, C., Votintseva, A. A., Walker, T. M., Cole, K., Davies, J., Fermont, J. M., Gascoyne-Binzi, D. M., Kohl, T. A., Kong, C., Lemaitre, N., Niemann, S., Paul, J., Rogers, T. R., Roycroft, E., Smith, E. G., Supply, P., Tang, P., Wilcox, M. H., Wordsworth, S., Wyllie, D., Xu, L., and Crook, D. W. (2016). Rapid, comprehensive, and affordable mycobacterial diagnosis with whole-genome sequencing: A prospective study. The Lancet Respiratory Medicine, 4(1):49–58.

PHE (2016). Tuberculosis in England: 2016. Technical report, Public Health England, London.

Renshaw, S. A., Mena-Allauca, M., Touriz, M., Renshaw, A., and Gould, E. W. (2014). The impact of template format on the completeness of surgical pathology reports. Archives of Pathology & Laboratory Medicine, 138(1):121–4.

Sedlmair, M., Meyer, M., and Munzner, T. (2012). Design study methodology: reflections from the trenches and the stacks. IEEE Transactions on Visualization and Computer Graphics, 18(12):2431–2440.

Shah, A. K. and Oppenheimer, D. M. (2008). Heuristics made easy: an effort-reduction framework. Psychol Bull, 134(2):207–222.

Shneiderman, B. (1996). The eyes have it: a task by data type taxonomy for information visualizations. Proceedings 1996 IEEE Symposium on Visual Languages.

Valenstein, P. N. (2008). Formatting pathology reports: applying four design principles to improve communication and patient safety. Archives of Pathology & Laboratory Medicine, 132(1):84–94.

Vredenburg, K., Mao, J.-Y., Smith, P. W., and Carey, T. (2002). A survey of user-centered design practice. Proceedings of the SIGCHI conference on Human factors in computing systems Changing our world, changing ourselves - CHI ’02, (1):471.

Walker, T. M., Kohl, T. A., Omar, S. V., Hedge, J., Del Ojo Elias, C., Bradley, P., Iqbal, Z., Feuerriegel, S., Niehaus, K. E., Wilson, D. J., Clifton, D. A., Kapatai, G., Ip, C. L. C., Bowden, R., Drobniewski, F. A., Allix-Bèguec, C., Gaudin, C., Parkhill, J., Diel, R., Supply, P., Crook, D. W., Smith, E. G., Walker, A. S., Ismail, N., Niemann, S., Peto, T. E. A., Davies, J., Crichton, C., Acharya, M., Madrid-Marquez, L., Eyre, D., Wyllie, D., Golubchik, T., and Munang, M. (2015). Whole-genome sequencing for prediction of Mycobacterium tuberculosis drug susceptibility and resistance: A retrospective cohort study. The Lancet Infectious Diseases, 15(10):1193–1202.

Wright, P., Jansen, C., and Wyatt, J. C. (1998). How to limit clinical errors in interpretation of data. Lancet, 352(9139):1539–1543.

Zipkin, D. A., Umscheid, C. A., Keating, N. L., Allen, E., Aung, K., Beyth, R., Kaatz, S., Mann, D. M., Sussman, J. B., Korenstein, D., Schardt, C., Nagi, A., Sloane, R., and Feldstein, D. A. (2014). Evidence-based risk communication: a systematic review. Annals of Internal Medicine, 161(4):270–80.

